# Conserved cis-acting motifs and localization of transcripts and proteins of *MFT2* in barley and rice

**DOI:** 10.1101/2023.01.17.524348

**Authors:** Shigeko Utsugi, Hiroyuki Kawahigashi, Akemi Tagiri, Rie Kikuchi, Kohei Mishina, Hiromi Morishige, Shingo Nakamura

## Abstract

*MFT* is an important regulator of seed dormancy in flowering plants. A natural mutation in the A-box motif in the promoter of wheat (*Triticum aestivum) MFT2* on chromosome 3A (*TaMFT2-3A*) has been used to prevent pre-harvest sprouting (PHS) in wheat cultivars in East Asia. Previous research using *in situ* hybridization showed that *TaMFT2-3A* is primarily expressed in the seed scutellum. In this study, we analyzed the localization of transcripts and encoded proteins for the *TaMFT2* homologs from barley (*Hordeum vulgare*) and rice (*Oryza sativa), HvMFT2* and *OsMFT2*, respectively. RNA *in situ* hybridization showed that, like the wheat genes, the rice and barley homologs are primarily expressed in the scutellum, indicating that these three *MFT2* genes have a common expression pattern during seed development. Analysis of the *cis*-acting regulatory elements of their promoter sequences showed that the three *MFT2* promoters share eight seed-specific *cis*-acting RY motifs, which are binding sites for B3-domain transcription factors of the AFL-B3 and VIVIPAROUS1/ABI3-LIKE (VAL) families. In addition, we detected tandemly repeated and partially overlapping A-box motifs in the promoters of *HvMFT2, TaMFT2-3B*, and *TaMFT2-3D*, possibly explaining why the natural allele of *TaMFT2-3A* has been employed in breeding. We generated transgenic rice plants expressing nuclear-localized green fluorescent protein (NLS-2xGFP) and OsMFT2-GFP under the control of a 3-kb fragment of the *OsMFT2* promoter. Immunohistochemical staining using anti-GFP antibodies mainly detected GFP in the scutellum and scutellar epithelium, which is an important tissue for initiating germination upon seed hydration. We confirmed these results by confocal microscopy of GFP fluorescence in seeds. Our results suggest that *MFT2* expression in rice, barley, and wheat might be regulated by a similar network of transcription factors through multi-RY motifs in the *MFT2* promoters, with possible roles in scutellar epithelium development.

## INTRODUCTION

Members of the MFT (MOTHER OF FLOWERING LOCUS T [FT] and TERMINAL FLOWER 1 [TFL1]) families belong to the phosphatidylethanolamine-binding protein (PEBP) family, which is widely distributed from prokaryotes such as *Escherichia coli* to eukaryotes such as humans (*Homo sapiens*) (Karlgren et al. 2011). In flowering plants, the PEBP family can be divided into the FT, TFL1, and MFT subfamilies. As mosses and ferns possess genes from the *MFT* clade but not from the *FT* or *TFL1* clades, *MFT* genes are thought to be the ancestral form of the two other clades (Hedman et al. 2009).

*MFT* was first discovered based on its homology with *FT* in Arabidopsis (*Arabidopsis thaliana*) (Yoo et al. 2004), and was thought to be involved in the regulation of flowering time, like *FT*. However, a subsequent functional analysis failed to provide any evidence for such a role (Kikuchi et al. 2009). In addition, *MFT* genes exhibit a seed-specific expression pattern, calling into question any involvement of *MFT* genes in flowering.

A detailed analysis of MFT function using mutants in Arabidopsis revealed that *MFT* is a negative regulator of abscisic acid (ABA) responses during germination (Xi et al. 2010). Moreover, transcriptome profiling with microarrays and the analysis of quantitative trait loci (QTL) for seed dormancy in wheat (*Triticum aestivum*) revealed that *TaMFT2-3A* is the causal gene underlying a major seed dormancy QTL (Nakamura et al. 2011). A natural mutation in a *cis*-acting A-box motif in the *TaMFT2-3A* promoter has been deployed as an important genetic resource to prevent pre-harvest sprouting (PHS) in wheat cultivars grown in monsoon climate areas of East Asia where the rainy season starts around the same time as the wheat harvest season (Nakamura et al. 2011 and 2015). Since then, a role for *MFT* homologs in regulating seed dormancy has been reported in various plant species including rice (*Oryza sativa*) (Song et al. 2020, Yoshida et al. 2022).

Rice has two *MFT* genes, *OsMFT1* (Os06g0498800, located on chromosome 6) and *OsMFT2* (Os01g0111600, located on chromosome 1) (Song et al. 2018 and 2020). *OsMFT2* is the ortholog of *TaMFT2-3A*. A study used genome editing to knock out this gene in rice, thus showing that *OsMFT2* is also involved in regulating seed dormancy. In addition, like Heading data 3a (Hd3a, the rice FT homolog), OsMFT2 was recently demonstrated to form a transcriptional regulatory module with the 14-3-3 family protein G-BOX FACTOR 14-3-3h (GF14h) and the basic leucine zipper (bZIP) transcription factor OREB1 to control temperature-dependent germination (Yoshida et al. 2022).

The underlying mechanisms of seed development and maturation have been intensively investigated in Arabidopsis. Indeed, seed development and maturation are controlled by a network of master transcription factors that includes LAFL (LEC1, ABI3, FUS3, and LEC2) proteins: two LEC1-type HAP3 family CCAAT-box binding factors (LEAFY COTYLEDON1 (LEC1) and LEC1-LIKE (L1L), and three B3 domain-containing transcription factors of the AFL-B3 family (ABSCISIC ACID INSENSITIVE3 (ABI3), FUSCA3 (FUS3) and LEC2). The phase transition from seed development and maturation to seed germination and vegetative development is itself regulated by VIVIPAROUS1 (Vp1)/ABI3-LIKE (VAL) proteins, which are also B3 domain-containing transcription factors that repress the LAFL network (To et al. 2006, Suzuki et al. 2007, Tsukagoshi et al. 2007, Santos-Mendoza et al. 2008, Jia et al. 2014, Lepiniec et al. 2018, Jo et al. 2019). The B3 domain of the AFL-B3 members mediates the activation of downstream genes by specifically recognizing the RY motif in their promoters.

Several orthologous LAFL genes in other species such as rice, barley (*Hordeum vulgare*), and wheat have been suggested to be involved in a similar regulatory cascade (Lepiniec et al. 2018). In addition, the rice VAL gene *GERMINATION DEFECTIVE 1* (*GD1*) was shown to be the causal gene in germination-defective rice mutants (Guo et al. 2013). The B3 domain of GD1 binds specifically to the RY motif to modulate the expression of its downstream targets. Thus, the RY motif is not only present in the promoters of seed-specific genes that encode seed storage proteins but is also a pivotal *cis*-acting element for controlling seed development and maturation and the phase transition.

FT and TFL1 were shown to be mobile proteins (Corbesier et al. 2007, Tamaki et al. 2007, Jaeger and Wigge 2007, Conti and Bradley 2007). PEBP family members exhibit an evolutionarily highly conserved protein structure (Karlgren et al. 2011), raising the possibility that MFT might also be a mobile protein, which can be deduced from a comparison of *MFT* transcript and MFT protein accumulation patterns in different tissues.

Scutellar epithelium tissue plays a pivotal role in germination (Fincher 1989, Kaneko et al. 2002). Upon seed hydration, scutellar epithelium tissue produces α-amylase and the plant hormone gibberellic acid (GA) (Kaneko et al. 2002). This newly synthesized GA is transported to aleurone cells, where it stimulates the biosynthesis of hydrolytic enzymes like α-amylase to degrade reserve nutrients like starch in endosperm. The degraded molecules are transported to the embryo to support shoot and root development during germination. Despite its importance for germination, however, previous *in situ* hybridization analysis using wheat seeds showed that *MFT* was expressed only in the scutellum and coleorhiza, but not in scutellar epithelial cells (Nakamura et al. 2011).

In this study, we investigated where *MFT2* transcript and MFT protein accumulate using *in situ* hybridization and transgenic rice plants expressing a nucleus-localized double green fluorescent protein (NLS-2xGFP) and an OsMFT2-GFP fusion under the control of a 3-kb *OsMFT2* promoter fragment. We demonstrate that wheat, barley, and rice *MFT2* have similar expression patterns, and their promoters show both common and distinct *cis*-acting elements. Moreover, we determined that NLS-2xGFP and OsMFT2-GFP localize in the scutellar epithelium. Our results suggest that rice, barley, and wheat *MFT2* expression might be regulated by similar LAFL network and VAL transcription factors through multi-RY motifs in their promoters and raise the possibility that *MFT2* genes may play roles in repressing germination through scutellar epithelial cells.

## MATERIALS AND METHODS

### Plant materials

The Japanese two-rowed malting barley (*Hordeum vulgare*) cultivar (cv.) ‘Haruna Nijo’ was grown in our experimental field in Tsukuba. The *japonica* rice (*Oryza sativa*) cv. ‘Nipponbare’ was grown at 28 ° C/24 ° C (day/night) under natural light from the beginning of May until the beginning of October (approximately 14-h light/10-h dark photoperiod).

### Genomic DNA Extraction, PCR, and Sequencing

Genomic DNA was isolated from leaves using a DNeasy Plant Mini Kit (Qiagen), following the manufacturer’s protocol. Alternatively, a simple DNA extraction method was followed, as detailed below. First, leaves were incubated at 50°C for more than a day; the resulting dried leaves were then ground to powder with a 3-mm glass bead placed in a tube with a Tissuelyser mill (Qiagen). The ground samples were incubated at 4°C in DNA extraction buffer (200 mM Tris-HCl pH 7.5, 250 mM NaCl, 0.5% [w/v] SDS, 25 mM EDTA) overnight. Genomic DNA was used as template for PCR using TaKaRa Ex Taq or PrimeSTAR GXL DNA polymerase (Takara Bio), according to the manufacturer’s protocol. The PCR conditions and primer sequences are described in Supplementary information and Table S1. The amplified fragments were purified using a QIAquick Gel Extraction Kit (Qiagen), then sequenced using a 3730xl-Avant DNA Analyzer (Applied Biosystems) and BigDye Terminator version 3.1 reagents (Thermo Fisher Scientific). The sequences were analyzed using Sequencher version 5.2.4 (Gene Codes Corporation), DNASIS Pro sequence analysis software (version 2.1; Hitachi Solutions), and GENTYX Ver.13 software (Genetyx).

### *In situ* hybridization (ISH)

ISH was carried out using longitudinal embryo sections of seeds from the barley cv. Haruna Nijo and the rice cv. Nipponbare. For barley samples, a 339-bp DNA fragment corresponding to the 3’ end of the coding sequence and 3’ untranslated region (3’ UTR) of *HvMFT2* (accession # AK375179) was amplified by PCR using the primers HvMFT2-F1 and HvMFT2-R1 (Table S1). For rice samples, a 214-bp DNA fragment corresponding to the 3’ UTR of *OsMFT2* (accession# NM_001372198) was amplified by PCR using the primers HindIIIOsMTF2-F1 and PstISoMTF2-R1 (Table S1).

The DNA fragments were cloned into the pTAC-2 vector (BioDynamics Laboratory) and positive clones were sequenced. The sequence-confirmed clones were linearized using *EcoR*I and *BamH*I restriction enzymes and then used as template to produce antisense and sense probes using T7 and SP6 RNA polymerase, respectively. ISH was conducted with a digoxigenin-labeled RNA probe and immunological detection following the method of Kouchi and Hata (1993), with modifications to improve the reactions as described by Komatsuda et al. (2007).

### Analysis of *cis*-acting motifs in the *MFT2* promoter regions

The *MFT2* promoter sequences were obtained from the EnsemblPlants database (Yates et al. 2022, https://plants.ensembl.org/index.html, rice cv. Nipponbare: IRGSP-1.0; barley cv. ‘Morex’: Morex version 3; wheat cv. ‘Chinese Spring’: IWGSC; *Arabidopsis thaliana* accession Columbia, TAIR10; *Triticum dicoccoides* [*T turgidum* subspecies *dicoccoides]*, WEWSeq_v.1.0; *Aegilops tauschii, Aet_v4.0*]). The *MFT2* promoter sequence of barley cv. ‘Haruna Nijo’ and the wild ancestral form of barley (*H. vulgare* ssp. *spontaneum*) accession ‘OUH602 (H602)’ was obtained from the Barley DB http://viewer.shigen.info/harunanijo/index.php (Sakkour et al. 2022) or http://viewer.shigen.info/barley (Sato et al. 2021), respectively. The locations of various *cis*-acting motifs in the *MFT2* promoter sequences were identified using GENTYX Ver.13 software (Genetyx).

### Vector construction

A genomic *OsMFT2* (Os01g0111600) fragment was amplified by PCR. The PCR product of 6,026 bp in length comprised a 2,897-bp promoter fragment, 153 bp of 5’ UTR, 2,557 bp of the coding region with all four exons, and 358 bp of 3’ UTR. This fragment was then used as template to amplify various regions of the *OsMFT2* genomic region by PCR. The nopaline synthase terminator (*NosT*) fragment was amplified by PCR using the pEGAD vector (accession # AF218816) as template. The sequence encoding synthetic GFP variant S65T (sGFP) was amplified by PCR from the pTH2 vector (Niwa et al. 2003). To assemble the appropriate DNA fragments, an In-Fusion HD Cloning Kit (Clontech) was used. The detailed procedure and list of PCR primers used for vector construction are provided in Supplementary information and Table S1.

### Rice transformation and preparation of seeds from transgenic rice plants

The final transformation vectors were introduced into Agrobacterium (*Agrobacterium tumefaciens*) strain EHA101. Rice cv. Nipponbare was used for Agrobacterium-mediated transformation as previously described (Toki, 1997). Plants regenerated from transformed callus (T0) were selected on solid Murashige and Skoog (MS) medium (Murashige and Skoog, 1962) containing 50 mg L^−1^ hygromycin at 27°C and grown under 16-h light/8-h dark conditions. Regenerated plants were grown in the greenhouse for further experiments.

Genomic DNA was extracted from the leaves of T0 plants and used as template for PCR. PCR conditions and primer sequences are described in the Supplementary Information and Table S1.

At heading, the selected rice plants were labeled with plastic tags. Rice seeds were harvested at 20 or 40 days after heading (DAH) and stored at 4°C until use.

### Immunohistochemistry

Rice seeds were fixed with 4% (w/v) paraformaldehyde, embedded in paraffin on CT-Pro20 (Genostaff) using G-Nox (Genostaff) as a less toxic organic solvent than xylene, and sectioned at 6 μm in thickness. Tissue sections were de-paraffinized with xylene and rehydrated through a graded ethanol series and phosphate-buffered saline (PBS). Antigen retrieval was performed by boiling for 10 min in a microwave, with a citrate buffer for heat-induced antigen retrieval (G-Activate buffer pH 6.0, Genostaff). Endogenous peroxidase activity was blocked by incubation with 0.3% (w/w) H_2_O_2_ in methanol for 30 min, followed by incubation with G-Block (Genostaff) and an avidin/biotin blocking kit (Vector Laboratories). The sections were incubated with 1 μg/ml anti-GFP rabbit polyclonal antibody (Thermo Fisher Scientific A11122) and normal rabbit Ig (Dako X0936) diluted with 1% G-Block (Genostaff)/TBS (Tris buffered Saline, 25 mM Tris, 137 mM NaCl, 2.68 mM KCl, pH 7.4) as negative control at 4°C overnight. The samples were washed two times with TBST (TBS with 0.1% Tween 20, pH 7.4) buffer for 5 min and one time with TBS buffer for 5 min at room temperature. The slides were then incubated with biotin-conjugated anti-rabbit Ig (Dako E0432) for 30 min at room temperature, followed by the addition of peroxidase-conjugated streptavidin (Nichirei Bioscience) for 5 min. Peroxidase activity was visualized by the addition of 3,3’-diaminobenzidine (DAB). The sections were counterstained with Mayer’s Hematoxylin (Muto Pure Chemical), dehydrated, and then mounted in malinol (Muto Pure Chemical).

### Sectioning and capturing GFP fluorescence images

The harvested transgenic rice plant seeds were analyzed using an SZX12 stereo microscope (OLYMPUS) attached with a GFP fluorescence light detection equipment to evaluate GFP fluorescence in embryos. Rice seeds were fixed with 4% (w/v) paraformaldehyde before being placed into ClearSee solution (Fujifilm Wako Pure Chemical Co.) to make them transparent (Kurihara et al. 2015). The cleared seeds were cut in half. The resulting half-excised rice seeds with embryo were placed in embedding molds with 5% (w/v) agar (Nacalai Tesque) at 50°C before being placed in a refrigerator at 4°C to solidify the agar. The hardened agar-embedded samples were glued to a plate in preparation for sectioning using Aron Alpha glue (TOAGOSEI). Longitudinal sections (thickness 100 μm) were then produced using a vibrating-blade microtome MicroSlicer DTK-100N (DOSAKA) and suspended in a drop of water on a glass slide. Fluorescence was excited with a 488-nm argon laser and emission was collected in the 500–560 nm range. For counterstaining, the samples were stained with propidium iodide (PI) (0.1 μg/ml) for overnight at 4°C, then washed them with PBS several times. PI stains the nucleus and cell membranes. The sections were analyzed using an LSM700, Imager.Z2 (Zeiss) or FV1000-D (OLYMPUS) confocal laser scanning microscopes.

## RESULTS

### *In situ* hybridization (ISH) in barley

The barley *MFT* gene used in this study was designated *HvMFT2* based on its rice ortholog *OsMFT2*, although this gene was previously reported as *HvMFT1* (accession # AB447466) (Kikuchi et al. 2009). A BLAST search using the EnsemblPlants database (Yates et al. 2022) indicated that the gene ID for *HvMFT2* is HORVU.MOREX.r3.3HG0219250 on chromosome 3H, based on the barley cv. ‘Morex’ reference genome (current version 3, Mascher et al. 2017). As *OsMFT2* is orthologous to *TaMFT2-3A, HvMFT2* is also an ortholog of *TaMFT2-3A*. The syntenic chromosomal locations of these *MFT2* genes supported the notion that these genes are true orthologs. A full-length complementary DNA (cDNA) clone for *HvMFT2* from the barley malting cv. “Haruna Nijo” was previously reported (NIASHv3087D21) (Matsumoto et al 2011). We designed a probe for ISH against *HvMFT2* based on this sequence.

ISH analysis using longitudinal sections of developing embryos of the barley cv. ‘Haruna Nijo’ at 14 and 21 days after heading (DAH) revealed that *HvMFT2* is primarily expressed in the scutellum, with constant or occasional weak expression in the coleorhiza, epiblast, shoot, mesocotyl, and cortex in the radicle (Fig. 1, Supplementary Fig. S1). This result was mostly consistent with a previous report of the expression pattern of the wheat ortholog *TaMFT2-3A* (Nakamura et al. 2011), except that we detected no ISH signals for *HvMFT2* transcripts in vascular bundles of barley embryos. In agreement with the *TaMFT2-3A* expression pattern observed in wheat seeds (Nakamura et al. 2011), we also detected no ISH signal in scutellar epithelial cells (Fig. 1, Supplementary Fig. S1).

**FIGURE 1.**
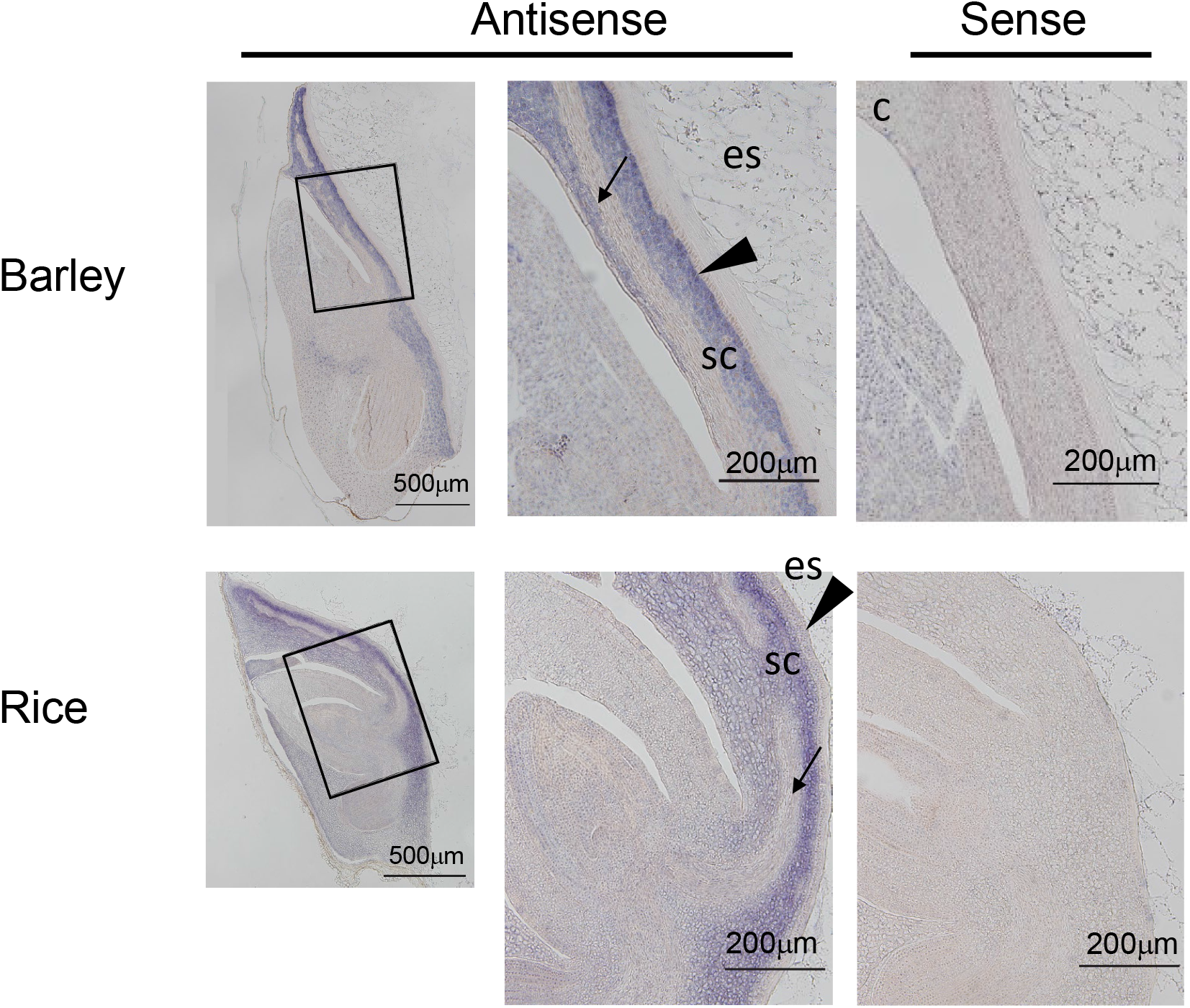
Localization of *MFT2* transcripts in immature barley and rice embryos. Longitudinal sections of immature embryos at 21 days after heading (DAH) from the barley cv. ‘Haruna Nijo’ and the rice cv. ‘Nipponbare’, respectively, probed with antisense or sense RNA. Positive hybridization signals are visible as a blue precipitate. es, endosperm; sc, scutellum. Triangles indicate the scutellar epithelium, and arrows indicate the vascular bundle.

### ISH in rice

We also carried out ISH using longitudinal sections of 14-DAH, 21-DAH, and mature (more than 40 DAH) seeds from the rice cv. ‘Nipponbare’ (NB). Similar to the results for barley *HvMFT2* above, we clearly observed *OsMFT2* expression in the scutellum, but not in vascular bundles of embryos or in scutellar epithelial cells, with constant or occasional weak expression in the coleorhiza, epiblast, mesocotyl, and cortex in the radicle (Fig. 1, Supplementary Fig. S2).

These results clearly underscored the highly similar tissue expression patterns of *HvMFT2* and *OsMFT2*. This expression pattern was also similar to that previously reported for *TaMFT2-3A* (Nakamura et al. 2011).

### *Cis*-acting regulatory motifs in the *MFT2* promoters in rice, barley, and wheat

*MFT2* genes of rice, barley, and wheat showed similar expression patterns (Fig. 1, Supplementary Fig. S1 and S2), suggesting that they might share regulatory *cis*-acting motifs in their promoters. To explore this possibility, we looked for known transcription factor binding sites including the RY motif (CATGCA), CCAAT motif (CCAAT), A-box motif (TACGTA), and G-box motif (CACGTG) over ~3 kb of promoter sequences upstream of the ATG codon, except for barley, for which we only looked at 2.4 kb of the promoter sequence, due to the presence of an upstream gene at this location.

We determined that *OsMFT2, HvMFT2*, and *TaMFT2-3A* promoters all have eight RY motifs, whereas the *OsMFT1* promoter contained only two RY motifs, and the *AtMFT* only one (Fig. 2). LEC1 and L1L in the LAFL network are CCAAT-binding transcription factors. As shown in Fig. 2, all promoters studied here have at least one CCAAT motif in their promoters, suggesting that LEC1 and L1L may participate in the regulation of *MFT2* transcription. A-box and G-box motifs are preferentially bound by bZIP transcription factors (Iazawa et al. 1993). The G-box is the core motif of the ABA-responsive element (ABRE). Our motif analysis identified an ABRE approximately 700 bp upstream from the ATG start codon in the *AtMFT* promoter (Fig. 2) as described by Xi et al. (2010). The ABRE consists of an RY motif and a G-box motif. The *OsMFT2* and *HvMFT2* promoters both had a G-box 284 bp and 305 bp upstream from the ATG start codon, respectively (Fig. 2), but lacked an adjacent RY motif to form an ABRE. In addition, we did not detect any G-box motif in the *TaMFT2-3A* promoter sequence (Fig. 2).

**FIGURE 2.**
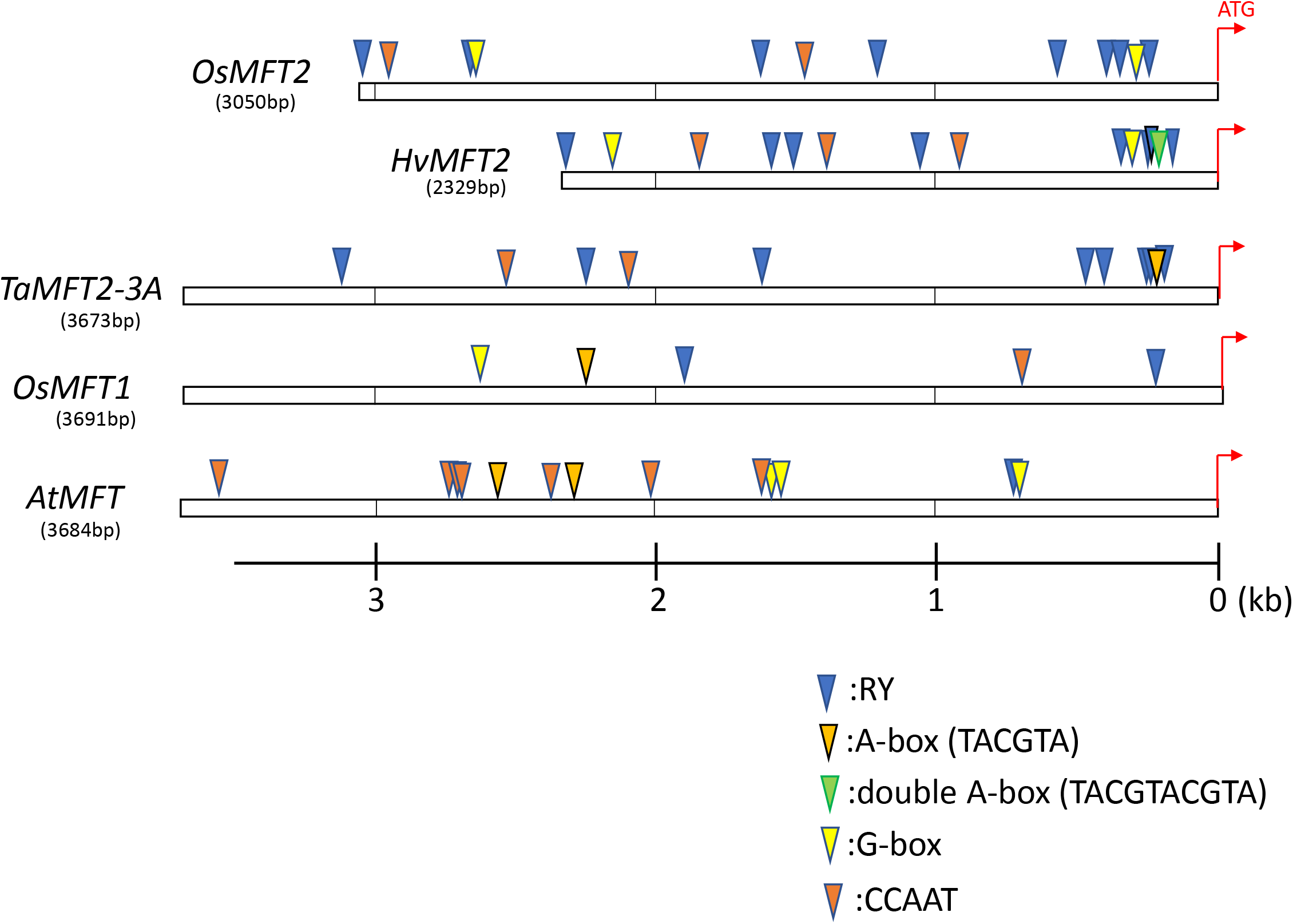
Localization of *cis*-acting motifs in *MFT2* promoter sequences from various plant species. Blue triangles, RY motif; orange triangles, A-box motif; green triangles, double A-box motif; yellow triangles, G-box; red brown triangles, CCAAT box. ATG, initiation codon of *MFT2* genes. Os, rice; Hv, barley; Ta, wheat; At, Arabidopsis.

We identified an A-box motif in the *TaMFT2-3A* and *HvMFT2* promoter sequences approximately 200 bp upstream from the ATG start codons but detected none in the *OsMFT2* promoter (Fig. 2). However, the *OsMFT1* and *AtMFT* promoter harbored one and two A-box motifs, respectively (Fig. 2), although they were located more than 2 kb upstream from the ATG start codons. We noticed that two A-box motifs are repeated in tandem with a 2-bp overlap (TACGTACGTA) in the *HvMFT2* promoter of the cv. ‘Haruna Nijo’. We designated this sequence as double A-box motif. We identified the same double A-box motif in the *MFT2* promoter of the conspecific wild barley ancestor (*Hordeum vulgare* ssp. *spontaneum*) accession ‘H602’ as well as in the cultivated barley cv. ‘Morex’. The bread wheat genome carries three other *MFT2* homologs, with two (*TaMFT2-3B-1* and *TaMFT2-3B-2*) on chromosome 3B and one (*TaMFT2-3D*) on chromosome 3D (Nakamura et al. 2011). Thus, we analyzed their promoter sequences for the A-box motif. They all had the same double A-box motif as in barley.

Our BLAST search using EnsemblPlants identified two *MFT2* orthologs (TRIDC3AG002240, *TdMFT2-3A* and TRIDC3BG001240, *TdMFT2-3B*) on chromosome 3A and 3B in *Triticum dicoccoides* (wild emmer wheat, tetraploid). These two genes appear to correspond to *TaMFT2-3A* and *TaMFT2-3B-2* in bread wheat, respectively. *TdMFT2-3A* and *TdMFT2-3B* also had the A-box motif and the double A-box motif in their promoters, respectively. Our BLAST search also detected one *MFT2* ortholog in Tausch’s goatgrass (*Aegilops tauschii*, AET3Gv20014800) on chromosome 3D, whose promoter had the double A-box motif 215 bp upstream from an ATG codon in the predicted exon I sequence which ends 159 bp upstream from the ATG codon.

### Localization of NLS-2xGFP in rice embryos

To better delineate the expression pattern of *OsMFT2*, we expressed *NLS-2xGFP* (encoding a nucleus-localized double GFP) under the control of a 3-kb *OsMFT2* promoter fragment (Fig. 3a). NLS-2xGFP itself has a very low transport rate through the nuclear pore complex (NPC) (Yang et al. 2004) and makes an excellent marker for the nucleus, like the previously used marker NLS-2xmOrange (Purwestri et al. 2009). We produced longitudinal sections of embryos from mature transgenic (T2 generation) seeds and probed them with an anti-GFP polyclonal antibody to detect the accumulation of GFP as a proxy for *OsMFT2* promoter activity. As shown in Fig. 4, we detected positive immunochemical staining signals in round-shaped nuclei as a brown color. This showed that NLS-2xGFP was specifically localized in the nucleus, as expected. Moreover, the signals were clearly visible in cells of the scutellum, as well as in the scutellar epithelium. In addition, we detected weak signals in cells of the plumule (coleoptile and first to third leaves) and the mesocotyl. However, we detected no signal around the shoot apex (Fig. 4).

**FIGURE 3.**
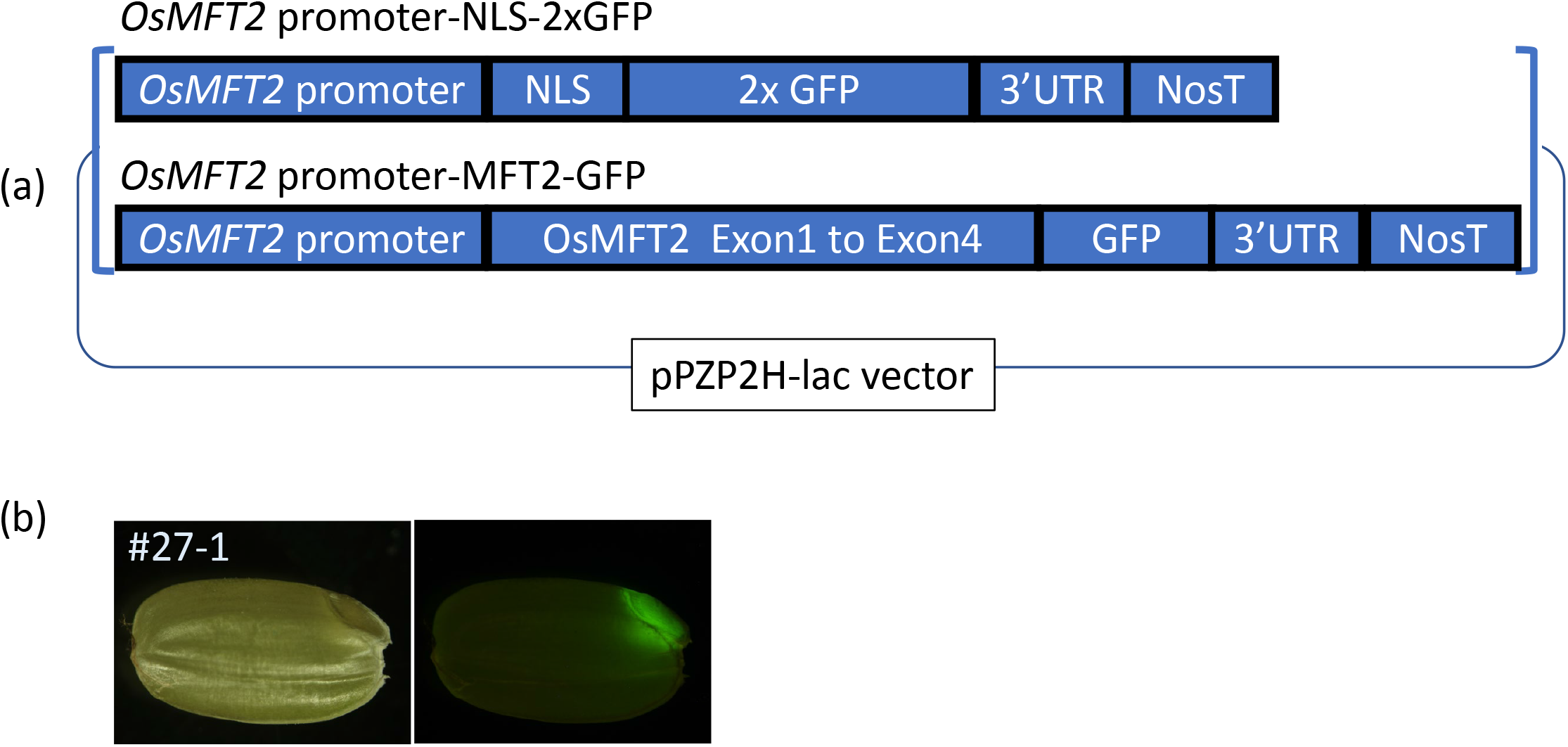
Expression domain of *OsMFT2* in rice seeds. (a) Schematic diagram of the constructs introduced into the rice cv. ‘Nipponbare’: *OsMFT2pro:NLS-2xGFP* and *OsMFT2pro:MFT2-GFP*. (b) Detection of GFP fluorescence in mature seeds (40 DAH) of T1 transgenic rice plant #27-1. Left, bright field view; Right, GFP fluorescence.

**FIGURE 4.**
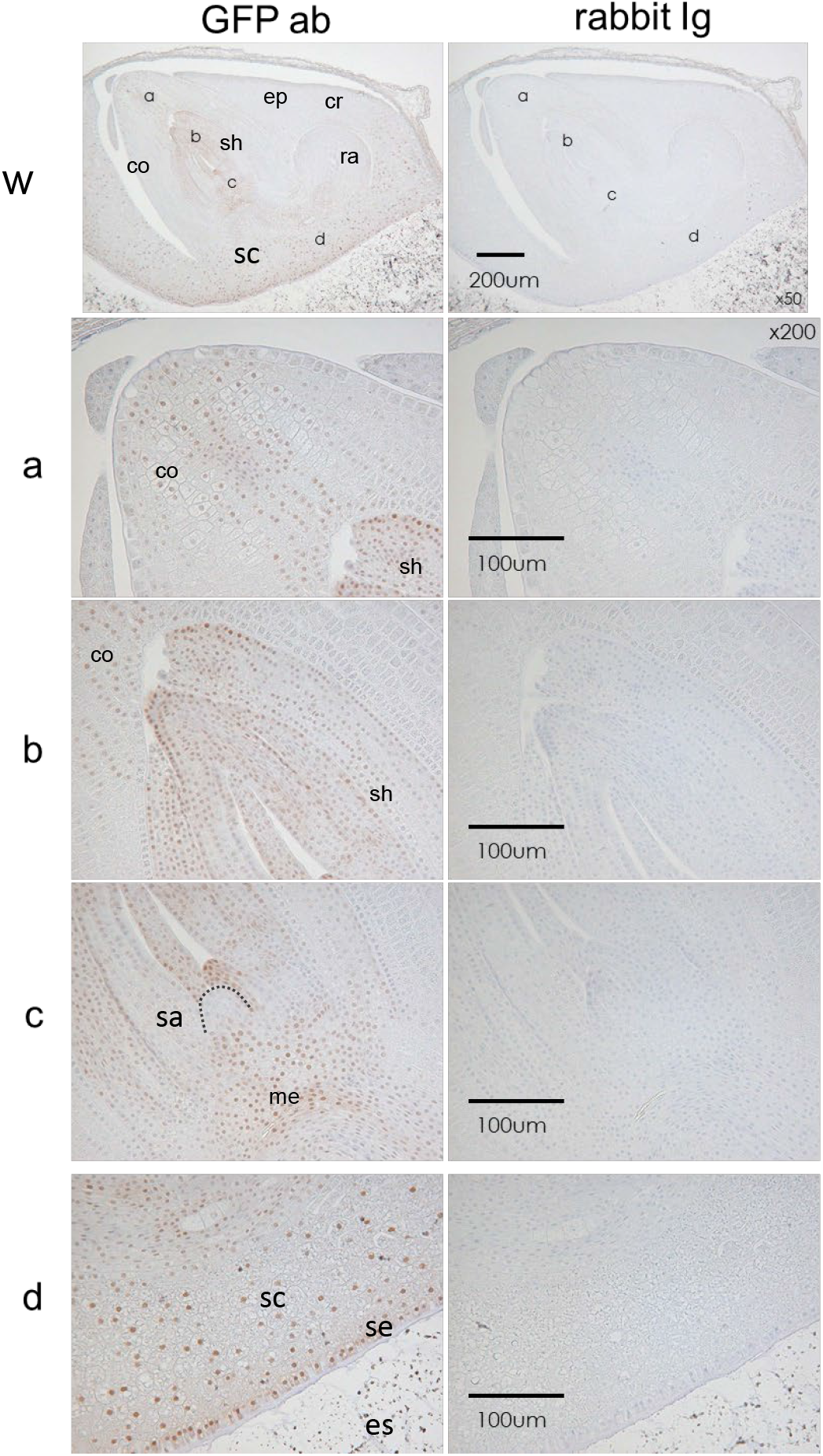
Localization of NLS-2xGFP protein in mature rice embryos by immunohistochemistry. Longitudinal sections of mature embryos at 40 DAH from rice transgenic plant #21-1. NLS-2xGFP was detected by immunohistochemical staining using an anti-GFP antibody (GFP ab) or rabbit IgG antibody (rabbit Ig) as negative control. w, whole embryo images. The images in a-c and d are magnified views of the areas indicated in w. The presence of GFP is indicated by the brown precipitates. Light blue color represents counterstaining with Mayer’s hematoxylin. co, coleoptile; cr, coleorhiza; ep, epiblast; es, endosperm; me, mesocotyl; ra, radicle; sa, shoot apex (surrounded by a dotted line); sc, scutellum; se, scutellar epithelium; sh, shoot (plumule); vb, vascular bundle.

### Localization of GFP-tagged OsMFT2

To analyze the subcellular localization and accumulation pattern of OsMFT2 in rice embryos, we generated another construct expressing *OsMFT2-GFP* (encoding OsMFT2 fused to GFP) under the control of a ~3-kb *OsMFT2* promoter fragment (Fig. 3a and b). The seeds from the transgenic rice lines showed strong GFP florescence in the scutellum in the embryos (Fig. 3b).

GFP was previously successfully employed as a fluorescent tag to track the movement of mobile FT and Hda3 from leaves to the shoot apical meristem in Arabidopsis and rice, respectively (Corbesier et al. 2007, Tamaki et al. 2007). We thus hoped to observe the movement of OsMFT2 with the GFP tag. As with the *OsMFT2pro:NLS-2xGFP* transgenic plants, we produced longitudinal sections of embryos from mature T2 40-DAH seeds of three independent lines (#2-1, #26-4, #27-1), using line #26-8 (having segregated the transgene out) as a negative control. We then probed these sections with an anti-GFP polyclonal antibody. As shown in Fig. 5, we detected clear positive immunochemical staining signals in the scutellum and scutellar epithelium, with constant or occasional weak signals in the coleorhiza, epiblast, coleoptile, mesocotyl and cortex in the radicle. On the subcellular level, we observed signals in nuclei and in the cytoplasm. Immunohistochemistry and *in situ* hybridization both failed to detect a signal in vascular bundle tissues of embryos.

**FIGURE 5.**
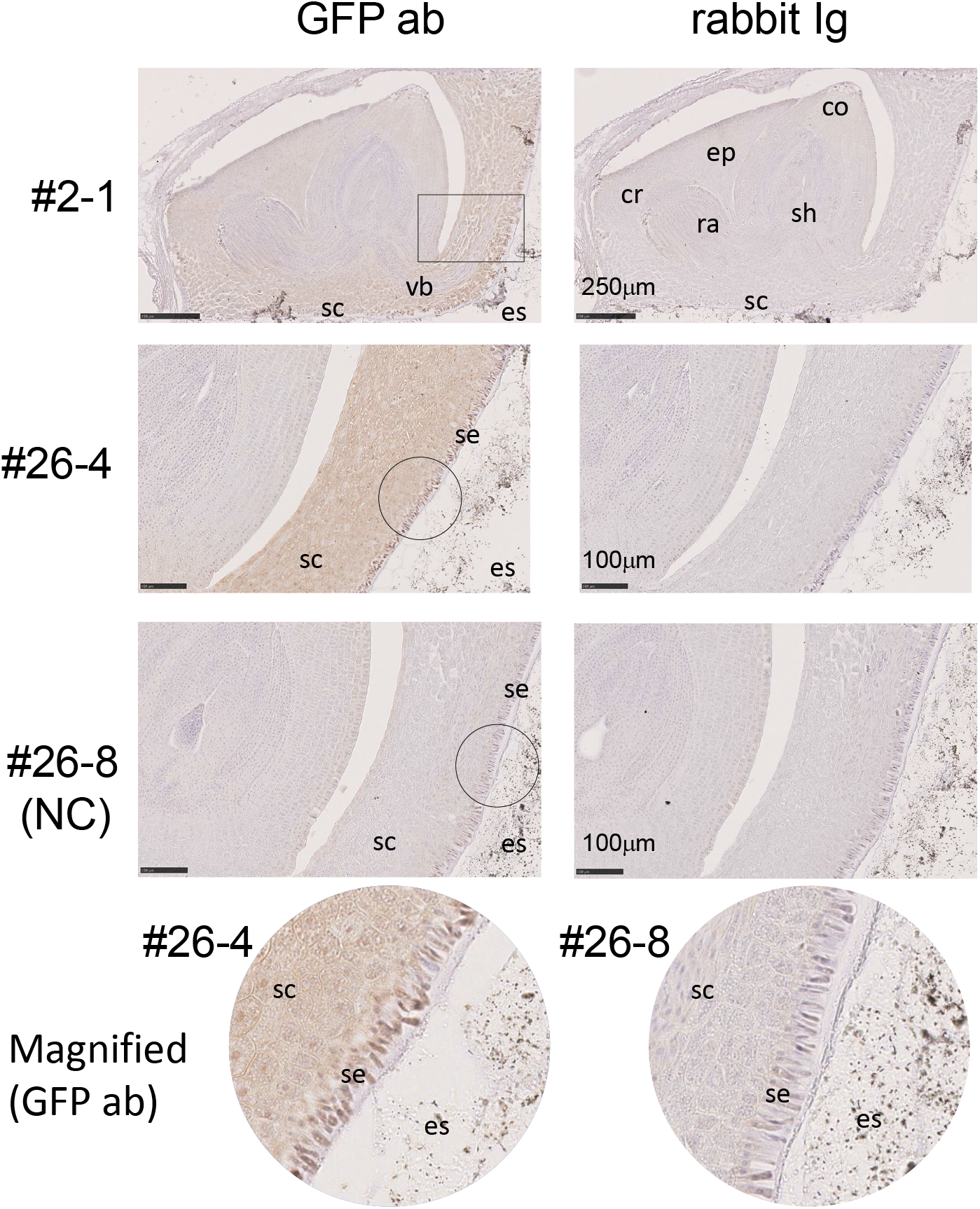
Localization of OsMFT2-GFP in mature rice embryos by immunohistochemistry. Longitudinal sections of the mature embryos at 40 DAH from T1 rice transgenic plants #2-1, #26-4, with plant #26-8 serving as a negative control (NC). OsMFT2-GFP was detected by immunohistochemical staining using an anti-GFP antibody (GFP ab) or rabbit IgG antibody (rabbit Ig) as negative control. #2-1: whole embryo images; #26-4 and #26-8, images of embryos from plants #26-4 and #26-8 showing a region equivalent to that highlighted by the rectangle for plant #2-1. Bottom, circular images are magnified views around the scutellar epithelial cell layers from the circles for plants #26-4 and #26-8. The presence of GFP is indicated by the brown precipitates. Light blue color represents counterstaining with Mayer’s hematoxylin. co, coleoptile; cr, coleorhiza; ep, epiblast; es, endosperm; me, mesocotyl; ra, radicle; sa, shoot apex (surrounded by a dotted line); sc, scutellum; se, scutellar epithelium; sh, shoot (plumule); vb, vascular bundle.

We further observed transgenic rice seeds at 20DAI by laser confocal microscopy. We visualized GFP fluorescence in the scutellum, scutellar epithelium, mesocotyl, coleorhiza, epiblast in embryos but not in the vascular bundle (Fig. 6). The GFP signal appeared to be present in both the nuclei and the cytoplasm. This result was consistent with the results obtained by immunohistochemistry.

**FIGURE 6.**
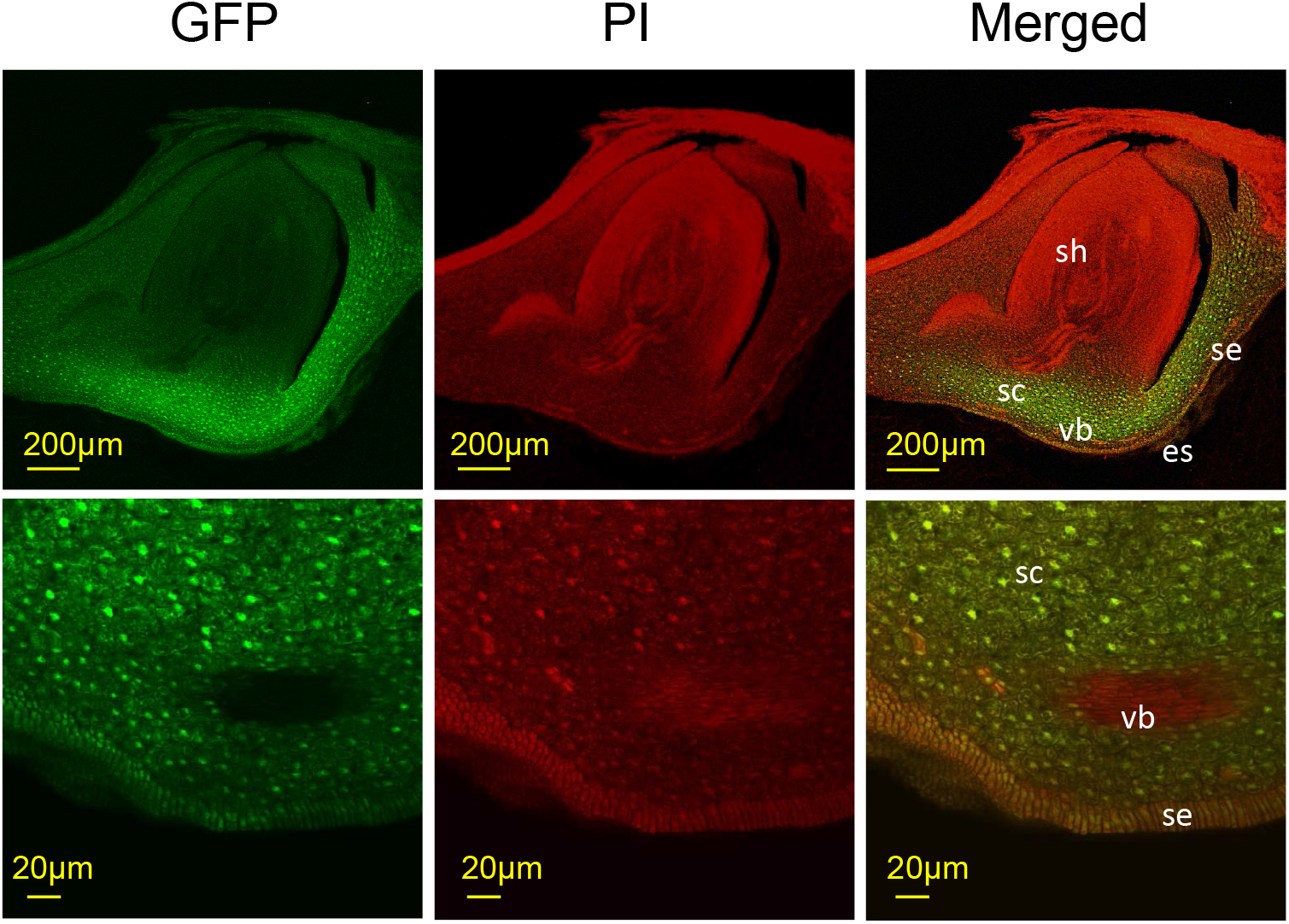
Localization of OsMFT2-GFP in mature rice embryos by confocal microscopy. Transverse sections of mature embryos at 40 DAH for T1 rice transgenic plant #27-1. MFT-GFP fluorescence was observed with a confocal laser scanning microscope. GFP, GFP fluorescent images (green color); PI, propidium iodide (PI) fluorescent images (red color); merged, merged images of GFP and PI fluorescence. co, coleoptile; cr, coleorhiza; ep, epiblast; es, endosperm; me, mesocotyl; ra, radicle; sa, shoot apex (surrounded by a dotted line); sc, scutellum; se, scutellar epithelium; sh, shoot (plumule); vb, vascular bundle.

## DISCUSSION

Based on the results of *in situ* hybridization experiments in this study and those from an earlier publication (Nakamura et al. 2011), the *MFT2* expression patterns appear to be highly similar among rice, barley, and wheat. These species belong to the *Poaceae* family and have albuminous seeds with a similar structure. These results suggest a functional similarity across *MFT2* orthologs between these species. In fact, in wheat and rice, the functional analysis of *MFT2* using transient assays and genome editing clearly showed that *MFT2* participated in the regulation of seed dormancy or germination (Nakamura et al. 2011, Song et al. 2020). Therefore, although we do not have equivalent functional evidence for *HvMFT2*, we propose that barley *MFT2* should also be involved in the regulation of seed dormancy.

Our results showed a weak signal for *MFT2* transcripts in coleorhiza, similar to the previous report on *TaMFT2-3A*. In barley, the coleorhiza was proposed to play a major role in regulating dormancy and germination by acting a barrier to the enclosed seminal root emergence through regulating ABA metabolism and sensitivity in this tissue (Barrero et al. 2009). *MFT2* genes in Arabidopsis and rice were previously shown to be involved in the regulation of germination through ABA sensitivity (Xi et al. 2010, Song et al. 2020). *MFT2* expression in the coleorhiza might contribute to regulating ABA sensitivity of this tissue to control germination.

To explore the common expression pattern of *MFT2* orthologs across species, we looked for the *cis*-elements in their promoters. Among the *cis*-regulatory motifs we investigated, the A-box motif and the RY motif showed distinct intriguing features. We determined that barley cv. ‘Haruna Nijo’ and its conspecific wild ancestor accession ‘H602’ harbored a double A-box motif (Fig. 2). This observation indicates that the double A-box is a common motif in the *MFT2* promoter in barley.

In barley, natural mutations in the A-box motif have not been reported, unlike in wheat. The double A-box motif in barley may have made it more difficult to select for natural mutations in this motif, leading to a more dormant phenotype. This result might also in part explain why we could not find barley cultivars with mutations in the A-box motif to improve PHS tolerance. Targeting the double A-box motif in barley, especially in the 2-bp overlapping sequence between the two A-box motifs, by genome editing may help generate a new useful allele for PHS tolerance in barley.

In bread wheat, the causal nucleotide substitution for the seed dormancy QTL, the first nucleotide (T) in the A-box motif of *TaMFT2-3A* is replaced by a C; this substitution is thought to inhibit decreases in the expression of *TaMFT2-3A* after the physiological maturity stage of seed development. This polymorphism makes seeds more dormant (Nakamura et al. 2011).

Bread wheat is derived via allopolyploid speciation through interspecific crossing between cultivated tetraploid wheat *Triticum turgidum* L. (AABB) and its diploid relative, *Ae. tauschii* Coss. (DD) (Miki et al. 2019). Cultivated tetraploid wheat was domesticated from the wild tetraploid wheat *T. dicoccoides* (AABB). The *MFT2* genes located on homoeologous chromosome A in the hexaploid and tetraploid wheat have the A-box motif, while the *MFT2* genes on homoeologous chromosome B and D in the hexaploid and tetraploid wheat have a double A-box motif. This genomic pattern raises the possibility that the A genome donor (*T. urartu*) may harbor the A-box motif, while the B genome donor candidate *Ae. speltoides* and the D genome donor *Ae. taushii* may carry the double A-box motif. Indeed, we identified the double A-box motif in the *MFT2* promoter from the *Ae.taushii* ortholog.

Among the four *MFT2* genes in hexaploid wheat, only *TaMFT2-3A* has the A-box motif. We speculate that this arrangement might be the major reason why mutations useful for PHS tolerance specifically affected *TaMFT2* on chromosome 3A, but not the *TaMFT2* genes on chromosome 3B or 3D.

The RY motif is one of the best studied *cis*-acting motifs involved in seed-specific gene expression such as seed storage protein genes (Fujiwara and Beachy 1994, Reidt et al. 2000). This motif is widely distributed in many seed-specific promoters of dicots and monocots (Reidt et al. 2000). In some cases, such as the *Napin* promoter, at least two RY motifs are redundantly involved in promoter activity (Reidt et al. 2000, Ellerström et al.1996, Stålberg et al. 1993). Perhaps some of the eight RY motifs detected in each *MFT2* promoter have redundant roles to regulate *MFT2* expression in rice, wheat, and barley.

The rice B3-domain transcription factor OsVP1 (rice ABI3 homolog) is one of the master regulators of the LAFL pathway in rice. *OsVP1* was reported to be expressed in vascular bundles, shoots and radicles (Miyoshi et al. 2002). These *OsVP1* expression patterns are clearly different from those observed for *OsMFT2* in this study, indicating that *OsMFT2* expression is unlikely to be directly regulated by OsVP1. The master regulators of LAFL and other transcription factors like bZIP transcription factors may instead coordinately regulate *OsMFT2* expression.

Due to their similarity in protein structure, MFT might be mobile, like FT and TFL1. However, we saw no evidence indicating that OsMFT2 is a mobile protein. However, our result cannot completely exclude this possibility.

Our immunohistochemistry results showed that OsMFT2-GFP localized both in the nuclei and the cytoplasm (Fig. 5). This is consistent with the previous results examining TaMFT2-3A-DsRed localization in onion leaf cells (Nakamura et al. 2011) and GFP-OsMFT2 in rice mesophyll protoplasts (Yoshida et al.2022). A recent study reported that the subcellular localization of OsMFT2 changed depending on its interaction with other proteins such as GF14h and OREB1 (Yoshida et al. 2022). Thus, our observation may reflect various protein-interaction states of OsMFT2.

We detected OsMFT2-GFP in scutellum epithelium cells by immunohistochemistry and by confocal microscopy. These results were contradictory to our *in situ* hybridization results. One possible explanation is that, due to technical limitations, the sensitivity of *in situ* hybridization might not be sufficient to detect *MFT2* transcripts in scutellar epithelial cells. Another possibility is that the repression of *OsMFT2* expression in the scutellar epithelium is controlled by cis motifs located outside of the *OsMFT2* sequence we used in this study. Our results using the *OsMFT2pro:GFP-OsMFT2* transgenic lines suggest that at least OsMFT2 can accumulate in scutellar epithelial cells in transgenic rice, which would be consistent with the critical regulatory functions of this cell type for the initiation of germination. This result might hold an important clue for future studies to understand the underlying functions of MFT2 in seed dormancy and germination for the important cereals.

### Future perspectives

We propose that *MFT2* in cereal crops may be directly controlled by the LAFL network and VAL proteins. A further detailed analysis of how the expression of *MFT2* is controlled and what kinds of downstream genes MFT2 regulates is necessary to understand how *MFT2* coordinately functions in gene regulatory networks for seed development, maturation and germination. Our results also suggest that MFT2 functions in scutellum epithelium cells in monocot cereal grains. This and previous results validate our interest in the roles of scutellum epithelium cells in terms of the regulatory mechanisms of germination or seed dormancy.

## AUTHORS AND CONTRIBUTIONS

SU, HK, AK, RK, KM, HM and SN performed the research and analyzed the data. HK and SN designed the research. SU, HK and SN wrote the article.

## ACKNOWLEDGMENTS

We deeply appreciate the advice for the rice transformation vector construction provided by Prof. K. Shimamoto (Nara Institute of Science and Technology). The sGFP vector was kindly provided by Prof. Niwa (University of Shizuoka). Dr. I. Ashikawa (Institute of Crop Science, NARO) kindly provided the pPZP2H-lac vector. Prof, K. Sato (Okayama University) kindly provided the *MFT2* promoter sequence for the barley cv. ‘Haruna Nijo’. Dr. T. Komatsuda at NIAS (current affiliation: Shandong Agriculture University in China) kindly permitted the use of his confocal laser scanning microscope LSM700. We are grateful to Genostaff Co. for their technical assistance for immunohistochemistry. ‘Haruna Nijo’ seeds were obtained through the National Bioresource Project of Barley, MEXT of Japan.

## SUPPLEMENTAL MATERIAL

Supplementary Material for this article can be found online at:

