## Supplementary information for "Conserved cis-acting motifs and localization of transcripts and proteins of *MFT2* in barley and rice"

**Detailed procedure for constructing the vectors for rice transformation**

A genomic *OsMFT2* (Os01g0111600) fragment was first amplified by PCR using the primers OsMFT2G-F1 and OsMFT2G-R2. Nested PCR was then carried out using the primers OsMFT2Pro-F1 and OsMFT2G-R1. The resulting amplicon was 6,026 bp in length and consisted of 2,897 bp of promoter, 153 bp of 5' untranslated region (5' UTR), 2,557 bp from exon 1 to exon 4, and 358 bp of 3' UTR. An In-Fusion HD Cloning Kit (Clontech) was used to assemble the DNA fragments.

1) *MFTpro:NLS-2xGFP* construction.

A sequence encoding three amino acids (M, A, P), followed by the nuclear localization signal (NLS) peptide from the simian virus (SV40) large T-antigen KKKRK, and one M residue was synthesized (5'-ATGGCTCCAAAGAAGAAGAGAAAGGTCATG-3'). The oligonucleotide encodes the protein sequence MAP-**KKKRK**V-M-, with the last M being the start codon of the first GFP. NLS-2xGFP has a very low transport rate through the nuclear pore complex (NPC) (Yang et al. 2004).

The nopaline synthase terminator (*NosT*) fragment was amplified by PCR using the cloning vector pEGAD (accession # AF218816) as a template, and the primers SalINosT-F1 and SphINosT-R1. The resulting *NosT* fragment was double digested with SalI and SphI restriction enzymes and cloned into the pHSG398 vector (Takara Bio) using a DNA Ligation Kit <Mighty Mix> (Takara Bio). The resulting clone was designated pHSG398NosT.

The sequence encoding the synthetic GFP S65T variant (sGFP) was amplified from the pTH2 vector (Niwa et al. 2003) as a template using the primers sGFP-F1 and sGFP-R1. To link the NLS sequence to the 5' end of the *sGFP* sequence, the *sGFP* fragment was amplified by PCR with the primers NLSGFP-F1 (containing the sequence encoding the NLS) and sGFP-R1. For TA-cloning (Thermo Fisher Scientific), a 3' adenine (A) overhang was added to both ends of the amplicons by incubation with Ex Taq polymerase (Takara Bio) at 72°C for 30 min in the presence of dATP before setting up the ligation reaction. Following the identification of positive clones and confirmation by sequencing, the cloned fragment was used as template for PCR using the primers EcoRINLSGFP-F1 and BamHIGFP-R1. The resulting amplicon was named EcoRI-NLS-GFP-BamHI and was double-digested with the EcoRI and BamHI restriction enzymes alongside the vector pHSG398NosT. Subsequently, the linearized vector and digested amplicon were ligated to yield the vector NLS-GFP-NosT/pHSG398 (clone #2 was selected).

The *sGFP* sequence was separately amplified with the primers BamHIGFP-F1 and

SalIGFP-R1. The resulting PCR product was A-tailed at both ends for cloning into the pCR4-TOPO vector using a Topo TA-cloning kit (Thermo Fisher Scientific), yielding the construct BamHI-GFP-SalI/pCR4-TOPO (clone #6 was used).

The constructs NLS-GFP-NosT/pHSG398 (clone #2) and BamHI-GFP-SalI/pCR4-TOPO (clone #6) were then digested with BamHI and SalI restriction enzymes. The GFP insert from BamHI-GFP-SalI/pCR4-TOPO was gel-purified and ligated into linearized NLS-GFP-NosT/pHSG398 using a Ligation Kit <Mighty Mix> (Takara Bio). Positive clones were selected for resistance to chloramphenicol (CM), yielding NLS-2xGFP-NosT/pHSG398 (clone #1 was used).

The vector pPZP2H-lac (Fuse et al. 2001) was digested with BamHI and HindIII restriction enzymes.

A fragment corresponding to the *MFT2* promoter and 5' UTR was amplified with the primers InFMFT2Pro-F1 and InFMFT2Pro-R1. The resulting amplicon was designated InFMFT2pro. The NLS-2xGFP-NosT region was amplified by PCR using NLS-2xGFP-NosT/pHSG398 (clone #1) as a template with the primers InFNLS-F1 and InFNosT-R1. Two bands were observed on 1% (w/v) agarose gel electrophoresis; the larger PCR product was excised from the gel and purified with a QIAquick Gel Extraction Kit (QIAGEN), resulting in a fragment designated InFNLS-2xGFP-NosT.

Finally, the BamHI- and HindIII-digested pPZP2H-lac, InFMFT2pro and InFNLS-2xGFP-NosT fragments were mixed and ligated together using an In-Fusion HD Kit. The resulting clone was designated MFT2ProNLSGFPx2NosT pPZPlac-2H (clone #4 was used).

2) *MFT-GFP* construction.

First, a 5,604-bp genomic fragment encompassing the promoter to exon 4 of *OsMFT2* was amplified by PCR using the primers InFMFT2Pro-F1 and InFMT2ProExon4-R1, resulting in the amplicon InFMFT2ProExon4.

The nopaline synthase terminator (*NosT*) fragment was amplified by PCR using the vector pEGAD (accession # AF218816) as a template, with the primers SalINosT-F1 and SphINosT-R1. The amplified *NosT* fragment was double digested with SalI and SphI restriction enzymes and cloned into pHSG398 vector (Takara Bio) using a DNA Ligation Kit <Mighty Mix> (Takara Bio). For In-Fusion cloning, the same fragment was amplified by PCR using pHSG398NosT as a template with the primers InFNosT-F1 and InFNosT-R1, yielding the amplicon InFNosT.

The sequence encoding synthetic GFP variant S65T (sGFP) was amplified from the pTH2 vector (Niwa et al. 2003) using the primers sGFP-F1 and -R1. The resulting sGFP amplicon was then used as a template for amplification by PCR with the primers InFGFP-F1 and InFGFP(PZP)-R1, producing InFGFP.

The 3' UTR fragment of *OsMFT2* for In-Fusion cloning was amplified by PCR using the nested PCR fragments of *OsMFT2* as a template with the primers InF3UR(PZP)-F1 and InF3UTR-R1 primers, resulting in the amplicon InF3UTR.

The pPZP2H-lac vector (Fuse et al. 2001) was double-digested with BamHI and HindIII restriction enzymes for In-Fusion cloning. The InFMFT2ProExon4 and InFGFP fragments and the digested pPZP2H-lac vector above were assembled by In-Fusion cloning, yielding MFT2ProExon4GFP-pPZP (clone #1 was used).

In parallel, the InF3UTR and InFNosT fragments were assembled into linearized pPZP2H-lac vector by In-Fusion cloning, producing 3'UTRNosT-pPZP (clone #1 was used). Positive clones were confirmed by Sanger sequencing.

For the final assembly, an amplicon was amplified by PCR from MFT2ProExon4GFP-pPZP (clone #1) as template with the primers InFMFT2Pro-F2 and InFGFP-R1. The resulting PCR product was named MFT2ProExon4GFP. Similarly, an amplicon was amplified by PCR using 3'UTRNosT-pPZP (clone #1) as a template with the primers InF3UTR-F1 and InFNosT-R2, producing 3'UTRNosT.

The MFT2ProExon4GFP and 3'UTRNosT fragments and the digested pPZP2H-lac vector above were assembled using an In-Fusion HD Cloning Kit.

In the resulting assembled clone, the order of elements was the *MFT* promoter, the 5' UTR, exon 1 to exon 4, *GFP* and *NosT*, inserted in the pPZP2H-lac vector. We designated this final clone MFT2ProExon4GFP3UTRNosTpPZP2H-lac, which was introduced into Agrobacterium (*Agrobacterium tumefaciens*) for Agrobacterium-mediated transformation of rice (Toki et al. 1997).

**Insert check for transgenic rice plants**

The presence of the T-DNA was checked by PCR, using genomic DNA extracted from leaves of $T_0$ plants using Takara Ex Taq DNA polymerase (Takara Bio) with the primers BamHIGFP-F1 and SalIGFP-R1 primers (*MFTpro:NLS-2xGFP* plants), or sGFP-F2 and OsMFT2-3UTR-R1 for (*MFT-GFP* plants). The PCR conditions were 35 cycles of 94°C for 20 sec, 60°C for 30 sec, 72°C for 1 min. The amplified fragments were checked by 1% (w/v) agarose gel electrophoresis.
