## Supplementary Figure S1 for "Conserved cis-acting motifs and localization of transcripts and proteins of *MFT2* in barley and rice"

Supplementary Figure S1| Localization of *MFT2* transcripts in embryos in barley detected by ISH

a) Immature embryos at 14 DAH, antisense probe    Bars: 200μm

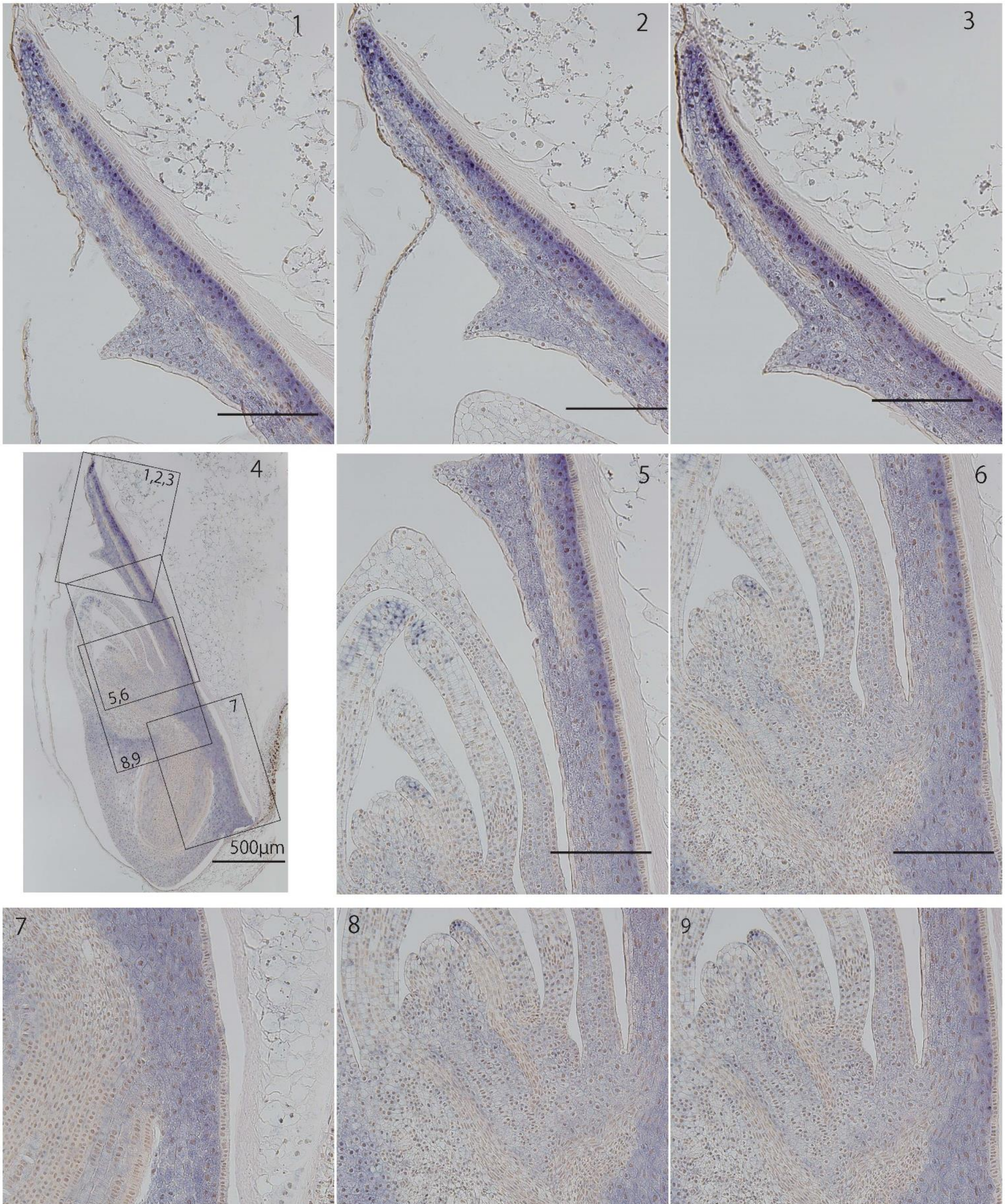

b) Immature embryos at 14 DAH, sense probe   Bars: 200μm

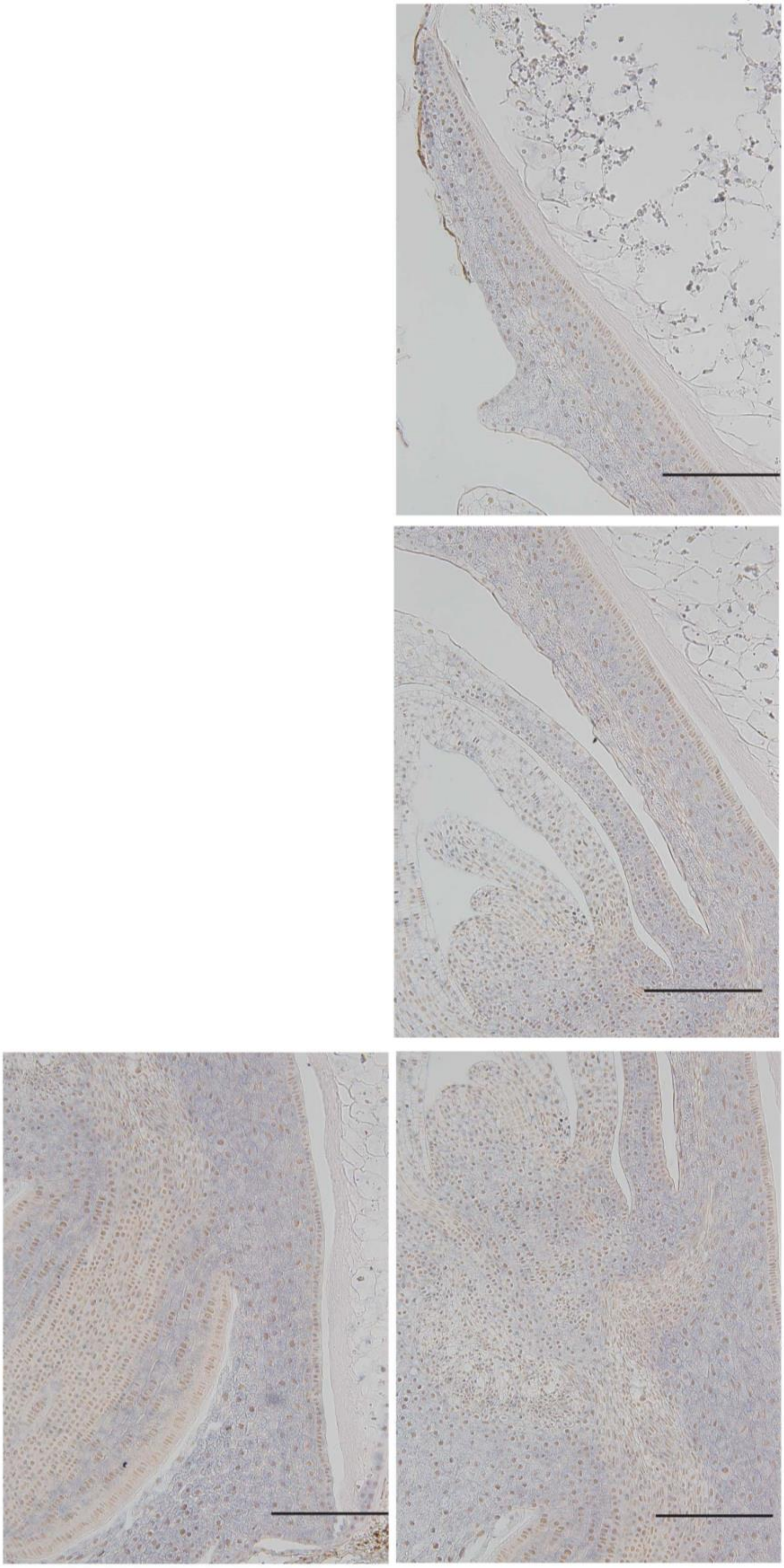

c) Immature embryos at 21 DAH, antisense probe    Bars: 200μm

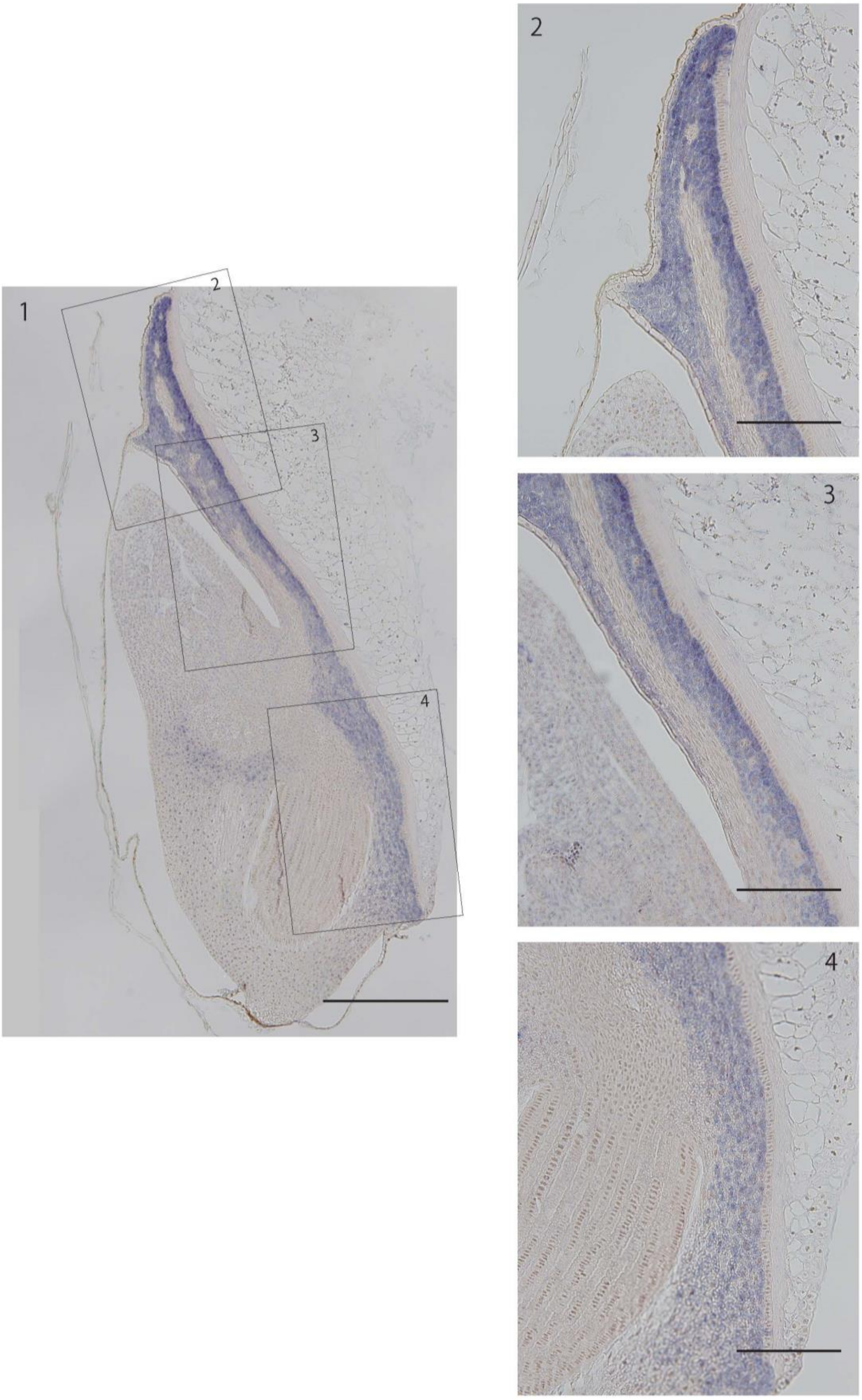

d) Immature embryos at 21 DAH, sense probe    Bars: 200μm

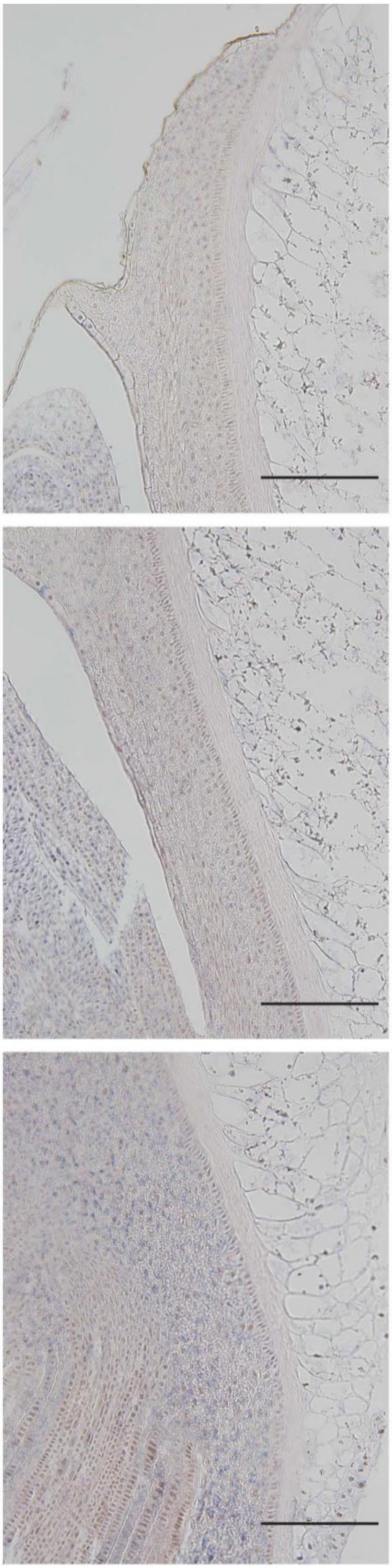
