## Supplementary Figure S2 for "Conserved cis-acting motifs and localization of transcripts and proteins of *MFT2* in barley and rice"

Supplementary Figure S2| Localization of *MFT2* transcripts in embryos in rice detected by ISH

a) Immature embryos at 14 DAH, antisense probe

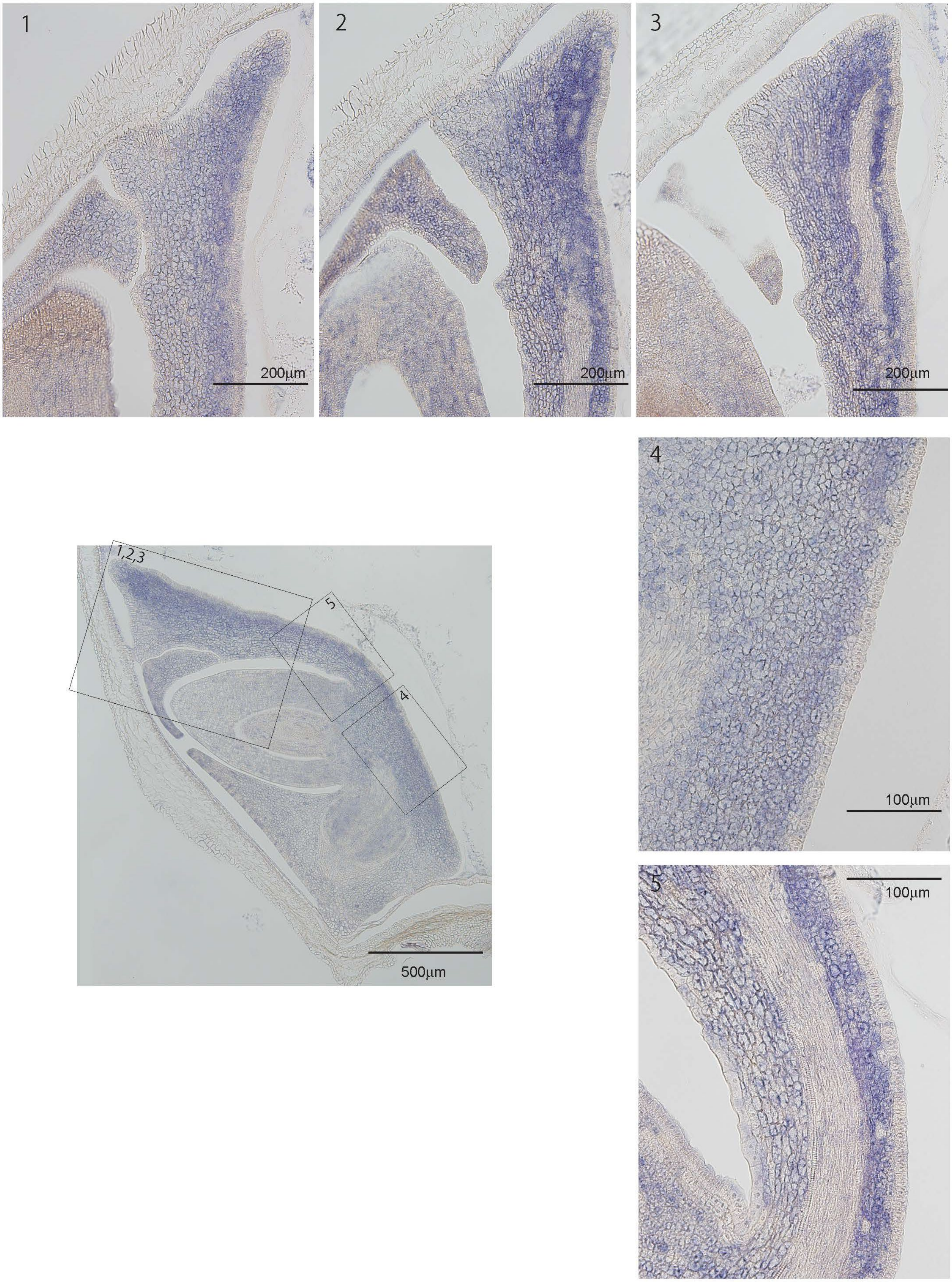

b) Immature embryos at 14 DAH, sense probe

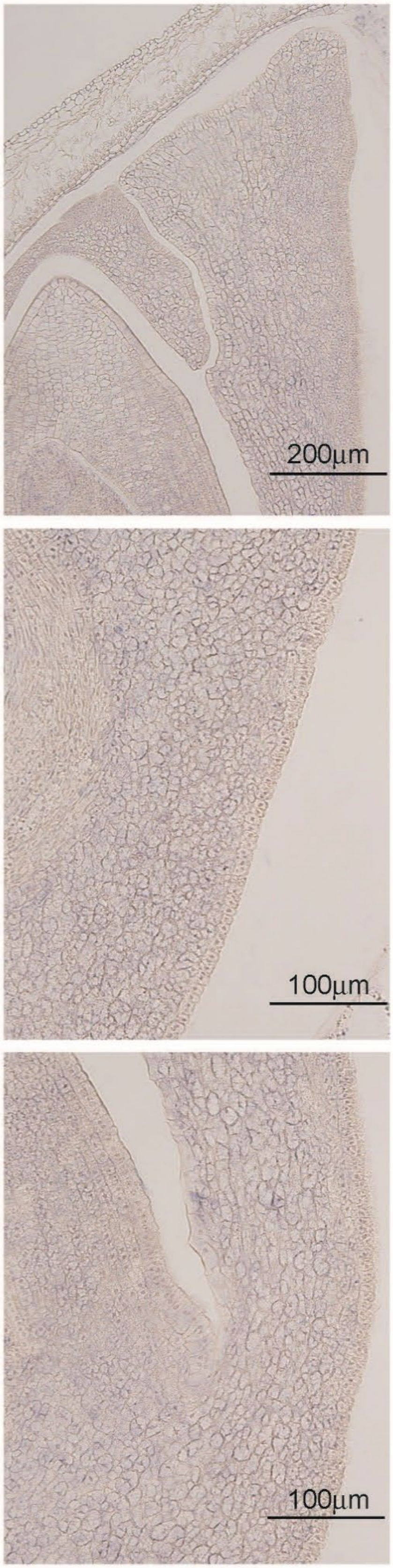

c) Immature embryos at 21 DAH, antisense probe    Bars: 200μm

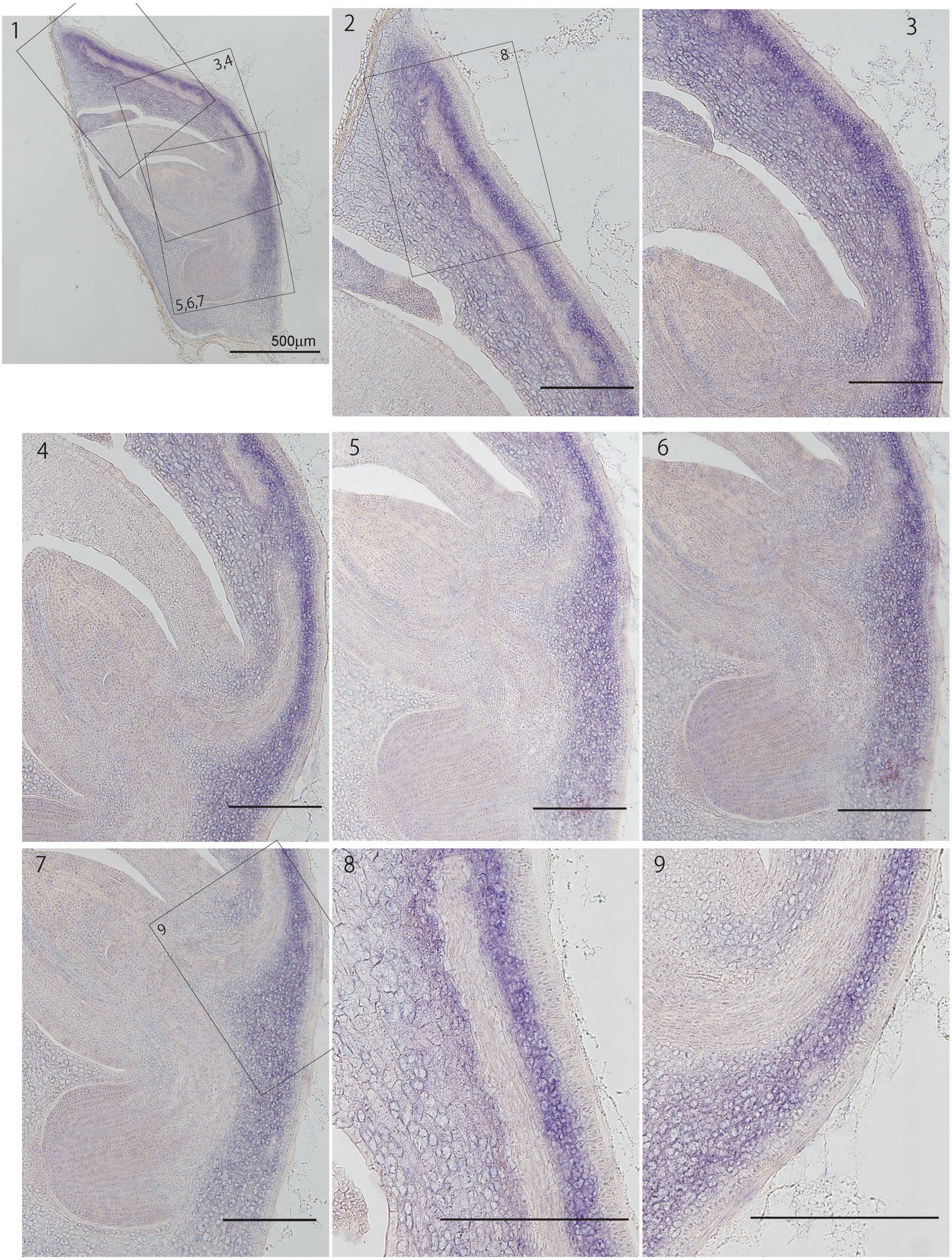

d) Immature embryos at 21 DAH, sense probe     Bars: 200μm

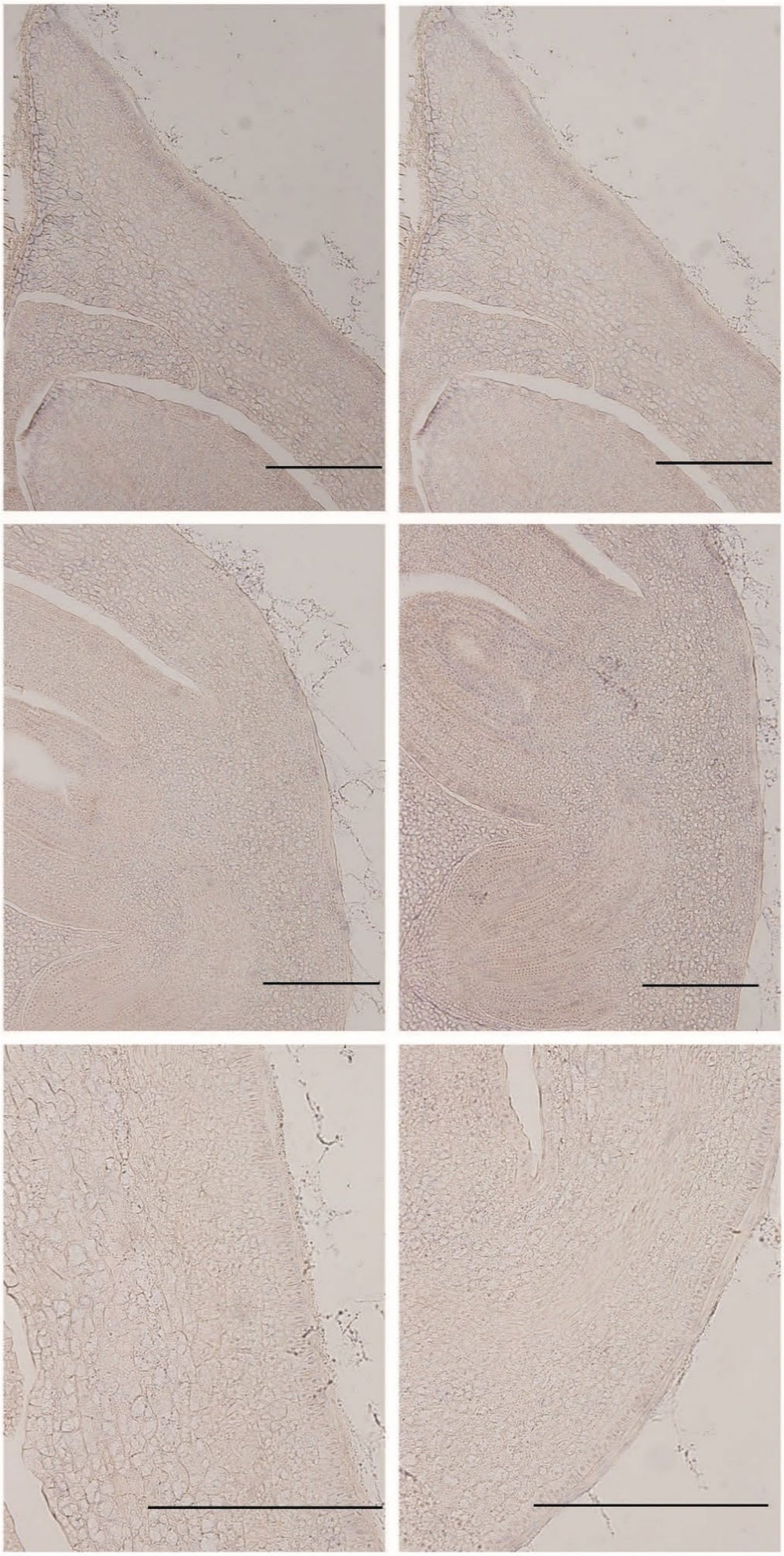

e) mature embryos at later than 40 DAH
antisense (a) and sense (s) probe

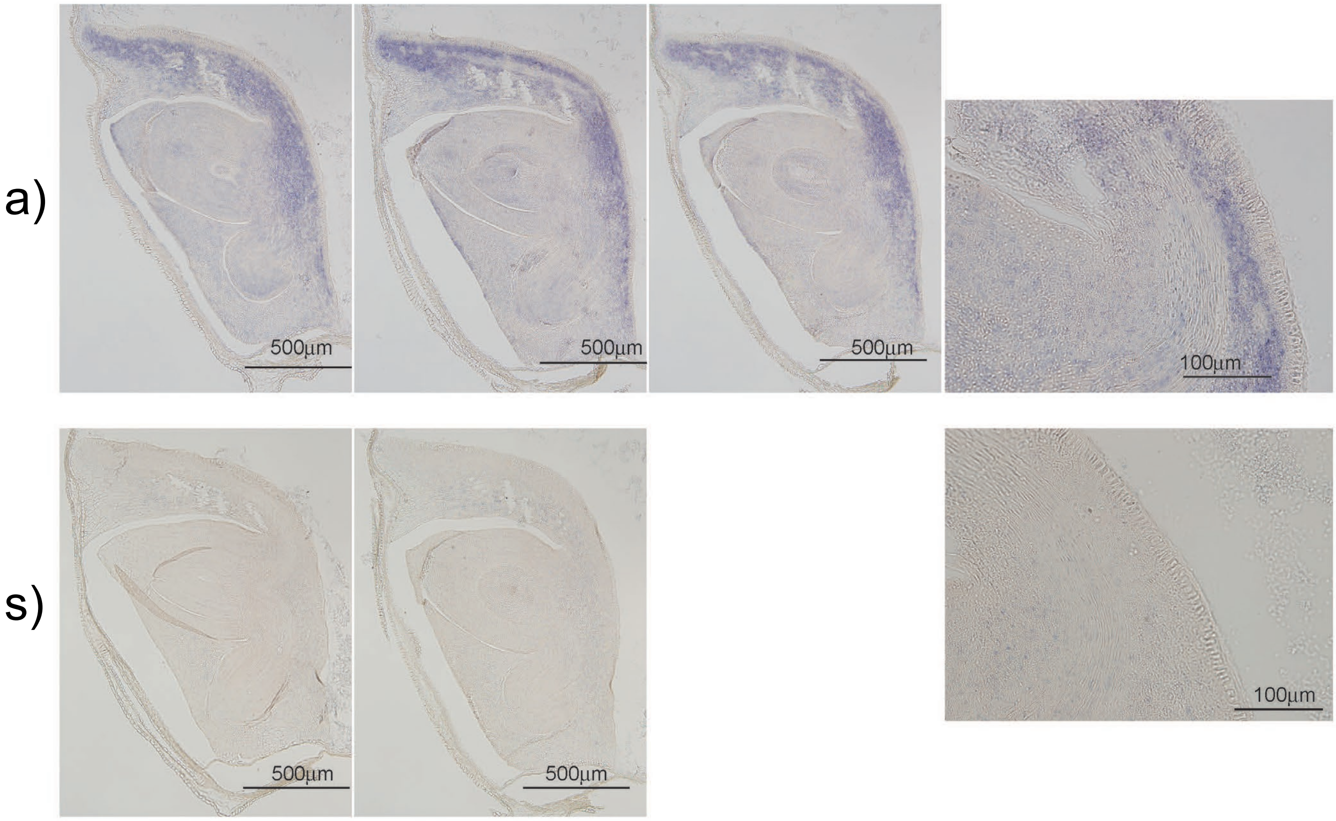
